# Serotype-dependent differences in AAV cellular transduction rates in the hypothalamus of Arctic ground squirrels

**DOI:** 10.64898/2026.05.13.724954

**Authors:** BW Laughlin, MH Sugiura, D Tupone, LE Fenno, MM Weltzin

## Abstract

Adeno-associated viral (AAV) vectors are foundational tools for dissecting brain structure–function relationships, but AAV serotype tropism varies across brain regions and species, requiring empirical validation to inform experimental design. This need is especially important in non-model organisms, where molecular neuroscience tools remain underdeveloped and access to research subjects is often limited. The Arctic ground squirrel (AGS, *Urocitellus parryii*) is a valuable model for studying extreme physiology, including metabolic suppression during hibernation and resistance to cerebral ischemia/reperfusion, yet no studies have evaluated AAV performance in the AGS brain. Here, we investigated the ability of AAV serotypes 1, 8, 9, and DJ to transduce the AGS hypothalamus using the human synapsin (hSyn) promoter and directly compared cellular transduction rates in a region implicated in thermoregulation and hibernation. To maximize data collection from a limited experimental population, we used a within-animal, contralateral stereotaxic injection design. Recombinant AAV vectors expressing enhanced green fluorescent protein or mCherry were delivered bilaterally, and reporter expression was analyzed four weeks later. All tested serotypes produced clear and reproducible reporter expression, establishing AAV as a viable molecular tool in the AGS hypothalamus. AAV1 produced significantly greater cellular transduction rates than AAV-DJ (17.2% ± 3.5% vs 8.4% ± 2.9%, paired *t*-test, *p* = 0.032). AAV8 and AAV9 showed transduction rates of 22.8% ± 0.6% and 20.1% ± 1.5%, respectively; however, with only two biological replicates per serotype, formal statistical comparison was not performed. These findings provide the first direct characterization of AAV-mediated gene delivery in the AGS brain and establish a foundation for future molecular interrogation of hypothalamic circuits in this extreme mammalian hibernator.

## Introduction

Studying non-model organisms enables investigation of specialized metabolic adaptations, relevant to brain energy metabolism, that are absent or attenuated in traditional laboratory models. In neuroscience, the mouse has served as the principal platform for the development of viral, genetic, and chemogenetic tools (Deisseroth 2015). Extending these approaches to non-model species requires direct evaluation of vector performance, as capsid tropism and transduction performance can vary across species and brain regions (Aschauer, Kreuz, and Rumpel 2013; Hordeaux et al. 2018).

Arctic ground squirrels (AGS) represent a valuable non-model system for studying extreme metabolic adaptations. During hibernation, AGS undergo profound metabolic plasticity including changes in the cerebral metabolic rate of glucose, and metabolic suppression. These changes are associated with neuronal plasticity and resistance to skeletal muscle and bone atrophy (Arendt and Bullmann 2013; Bogren et al. 2016; Drew et al. 2023; Goropashnaya et al. 2025; Richter et al. 2015). In addition, AGS exhibit tolerance to cerebral ischemia/reperfusion (I/R) injury (Bhowmick and Drew 2017; Dave et al. 2006). These adaptations provide a unique opportunity to investigate neuroprotective and metabolic mechanisms not readily accessible in traditional laboratory species.

Adeno-associated viruses (AAVs) are foundational tools in modern neuroscience, yet their performance must be validated in each biological context. AAV serotypes are capsid variants that differ in receptor interactions, tissue spread, and cellular tropism, resulting in context-dependent differences in transduction efficiency and relative bias toward neuronal or non-neuronal cell populations. Although AAV2 and AAV9 have been successfully used in 13-lined ground squirrels (Junkins et al. 2024; Zhang et al. 2021), serotype performance has not been systematically characterized in AGS. We therefore selected AAV1, AAV8, and AAV9 as commonly used naturally occurring capsids with established utility for neuronal gene delivery in rodent brain, together with AAV-DJ, a rationally engineered hybrid capsid reported to enhance transduction in some contexts. Because AGS used in this study were wild-caught rather than laboratory-bred, biological variability was expected and experimental replication was limited. Under these constraints, empirical validation of capsid performance in AGS brain tissue was essential to establish the feasibility and interpretability of future viral-mediated manipulations.

In this study, using adult Arctic ground squirrels, we directly compared the transduction rates of multiple AAV serotypes delivered by contralateral injections into the broader area surrounding the dorsomedial hypothalamus (DMH), a region implicated in thermoregulation and hypothalamic circuits relevant to hibernation and daily torpor (Ambler et al. 2022; Frare et al. 2019; Morrison et al. 2025; Morrison and Nakamura 2011).

## Methods

### Animals

All animal care and experimental procedures were conducted in accordance with the Guide for the Care and Use of Laboratory Animals and were approved by the Institutional Animal Care and Use Committee (IACUC) at the University of Alaska Fairbanks.

A total of eight wild adult AGS (*Urocitellus parryii*) were used in this study, including four females (616 ± 77 g) and four males (857 ± 150 g). Animals were captured during the summer of 2024, in the northern foothills of the Brooks Range, Alaska, approximately 40 miles south of Toolik Field Station (68°38′ N, 149°38′ W; elevation 809 m), and transported to the University of Alaska Fairbanks under an Alaska Department of Fish and Game permit.

### Husbandry

To deter entry into torpor, AGS were housed at an ambient temperature of 21 °C under a 12 h light/12 h dark photoperiod. Animals were singly housed in standard cages (12″ × 19″ × 12″) containing cotton nesting material, with food and water provided ad libitum throughout the study. Following stereotaxic injection, animals were maintained under these torpor-suppressing conditions for 4 weeks to allow for viral expression prior to tissue collection.

### Stereotaxic surgery

Animals were anesthetized with isoflurane (5% induction; 2–3% maintenance in oxygen at 1.0–1.5 L/min) and placed in a stereotaxic frame (Kopf Instruments, Tujunga, CA, USA). Ophthalmic ointment was applied to prevent corneal drying, and body temperature was maintained at 36–37 °C using a circulating water blanket. Heart rate, oxygen saturation, respiratory rate, and rectal temperature were monitored throughout the procedure.

Pre-operative analgesia consisted of extended-release meloxicam (4.0 mg/kg, subcutaneous), and sterile saline (2 mL, subcutaneous) was administered for hydration. The scalp was shaved and aseptically prepared using alternating scrubs of povidone–iodine and sterile water, followed by 70% isopropyl alcohol. Local anesthesia was provided using a subcutaneous lidocaine line block (0.5%, 5 mg/mL; ≤7 mg/kg) at the incision site.

A midline scalp incision was made to expose the skull, and bregma and lambda were identified and leveled in the horizontal plane (≤ 0.1 mm deviation). Craniotomies were performed using a dental drill at stereotaxic coordinates targeting the DMH (interaural +9.4 mm, ±0.5 mm lateral, −12.7 mm ventral). Anteroposterior coordinates were derived from the rat brain atlas using a 1.5× scaling factor obtained empirically from a comparative study in Tupone’s lab (data not published) to account for the larger brain size of AGS (Paxinos and Watson 2018).

Adeno-associated viral vectors (1,000 nL) were delivered via a pulled glass micropipette connected to a pressure-based microinjection system (PV830 Pneumatic PicoPump; World Precision Instruments, Sarasota, FL, USA) at a rate of 100 nL/min. To enable within-animal comparisons, distinct AAV serotypes with unique reporter genes were injected into the left and right DMH of the same animal. Following injection, the micropipette was left in place for 2 min to minimize reflux before slow withdrawal. The scalp was closed using sutures.

Following surgery, isoflurane was discontinued and animals were allowed to recover under supplemental oxygen and warming until ambulatory. Animals were monitored closely during recovery and daily thereafter. Enrofloxacin (5 mg/kg, subcutaneously) was administered for 5 days to prevent infection. Incision sites were inspected routinely, and sutures were removed 10–14 days postoperatively. Four weeks after viral delivery, animals were euthanized, and brains were extracted for histological analysis.

### Adeno-associated viral vectors

Recombinant adeno-associated viral (AAV) vectors encoding enhanced green fluorescent protein (EGFP) or mCherry under the control of the human synapsin promoter (hSyn) were used to assess transduction rates. Serotypes were selected to represent both commonly used naturally occurring capsids (AAV1, AAV8, AAV9) and a rationally engineered hybrid capsid (AAV-DJ) with reported enhanced transduction performance (Grimm et al. 2008).

AAV1 and AAV9 have demonstrated robust neuronal tropism in rodent central nervous system applications, AAV8 has been associated with high transduction performance across multiple tissues including brain, and AAV-DJ was included based on its engineered properties designed to improve infectivity and cellular entry (Aschauer et al. 2013).

Recombinant pAAV1-hSyn-EGFP (#50465-AAV1; 1.9 × 10^13^ genome copies [GC]/mL), pAAV9-hSyn-mCherry (#114472-AAV9; 2.4 × 10^13^ GC/mL), and pAAV8-hSyn-EGFP (#50465-AAV8; 3.6 × 10^13^ GC/mL) were obtained from Addgene (Watertown, MA, USA). AAV-DJ-hSyn-Cherry (#GVVC-AAV-017; 3.65 × 10^13^ GC/mL) was obtained from the Stanford University Gene Vector and Virus Core (Stanford University, Stanford, CA, USA). Prior to stereotaxic injection, viral stocks were diluted 1:10 in sterile 0.9% NaCl without normalization of genome copy concentration across serotypes. Viral stocks were diluted to a common injection dilution rather than normalized by genome copy concentration across serotypes, consistent with prior studies noting practical limitations in direct titer-matched comparisons of AAV vectors (Fenno et al. 2020). A total volume of 1,000 nL of virus was delivered per injection site. Animals were housed in ambient temperature of 21 °C under a 12 h light/12 h dark photoperiod to prevent hibernation for four weeks following viral delivery to permit stable transgene expression before tissue collection.

### AGS perfusion and Tissue processing

AGS were anesthetized with isoflurane (5% induction) and maintained at 3% via facemask with 100% medical-grade oxygen delivered at a flow rate of 1.5 L/min. Animals were transcardially perfused with 0.9% NaCl, followed by fixation with 4% formaldehyde in 10 mM PBS containing 0.9% NaCl. Brains were removed and post-fixed in 4% formaldehyde for 12 h at 4 °C, then cryoprotected in sucrose solutions of increasing concentration (5%, 15%, and 30%) until fully infiltrated. Coronal sections (40 µm) were cut using a CryoStar NX50 cryostat (Thermo Fisher Scientific, Waltham, MA, USA) and collected into four serial sets (every fourth section per set).

### Immunohistochemistry

Coronal brain sections were mounted onto glass slides and processed for immunohistochemistry. Sections were rinsed in phosphate-buffered saline (PBS; 3 × 5 min) and permeabilized in PBS containing 0.3% Triton X-100 for 15–30 min at room temperature. Non-specific binding was blocked for 1 h at room temperature in antibody dilution solution (ABS; sodium phosphate-buffered saline containing 0.3% Triton X-100, 0.9% NaCl, 0.25% λ-carrageenan, 0.02% sodium azide, and 1% normal donkey serum) with gentle agitation.

Mounted sections were incubated overnight at 4 °C with primary antibodies diluted in ABS: rabbit anti-mCherry (1:1000; Abcam, Cambridge, MA, USA; cat. #ab213511), chicken anti-GFP (1:1000; Abcam, Cambridge, MA, USA; cat. #ab13970), or rabbit anti-NeuN (1:2500; Invitrogen, Waltham, MA, USA; cat. #PA5-78499). Following primary antibody incubation, sections were washed in PBS (3 × 10 min) and incubated for 2 h at room temperature with species-appropriate Alexa Fluor secondary antibodies diluted 1:500 in ABS. Reporter-labeled sections received donkey anti-rabbit Alexa Fluor 594 (Cat. #A-21207) and donkey anti-chicken Alexa Fluor 488 (Cat. #A-78948), whereas NeuN-labeled sections received donkey anti-rabbit Alexa Fluor 647 (Cat. #A31573) (Thermo Fisher Scientific, Waltham, MA, USA). All incubations were performed with gentle rocking and protected from light.

After secondary antibody incubation, sections were washed in PBS (3 × 10 min), coverslipped with ProLong™ Gold Antifade Mountant with DAPI (Cat. #P36931; Thermo Fisher Scientific, Waltham, MA, USA), and allowed to cure overnight at room temperature in the dark prior to imaging.

### Image acquisition and data analysis

Fluorescence images were acquired using a Nikon Eclipse E80i epifluorescence microscope (Nikon Corporation, Tokyo, Japan) equipped with a high-sensitivity monochrome back-illuminated camera (BSI Express) and an X-Cite 120PC metal halide light source (Excelitas Technologies, Waltham, MA, USA). Image acquisition was performed using NIS-Elements AR software (version 5.30.06; Nikon). Low-magnification images (4×) were acquired and stitched to generate anatomical overviews of reporter expression within the hypothalamus. High-magnification images (40×) were acquired for quantitative analysis. For each coronal level, one to three non-overlapping 40× fields were collected per section, depending on the extent of reporter expression. Illumination intensity was held constant across imaging sessions, and exposure settings were adjusted as needed to avoid saturation while preserving reporter signal detection.

Animals were included in paired analyses only if detectable reporter expression was observed for both serotypes in the targeted hypothalamic region. One animal exhibiting reporter expression for only a single serotype and one animal lacking adequate reporter expression for both serotypes were excluded from paired statistical comparison. These exclusions were based on predefined criteria for within-animal comparison and do not distinguish between technical and biological sources of variability.

Quantitative image analysis was performed using Image-Pro (version 11.1; Media Cybernetics, Rockville, MD, USA). Scale calibration was verified prior to analysis, and all measurements were performed on 40× images. Composite images containing DAPI and reporter channels were generated for analysis. Nuclei were identified using an AI-based deep learning model applied to the DAPI channel (diameter parameter: 5.16; strictness: 35%; probability threshold: 50%). A ring-region protocol was applied to generate a perinuclear ring surrounding each nucleus to assess colocalization with reporter signal (grow = 5; offset = 1). This approach was selected to capture perinuclear cytoplasmic reporter signal while minimizing inclusion of adjacent neurites. Reporter signal was segmented using Smart Segmentation. Transduction rate was calculated for each 40× field as the proportion of DAPI-positive nuclei exhibiting colocalization with reporter signal (reporter-positive nuclei / total DAPI-positive nuclei × 100).

For neuronal colocalization analysis, separate NeuN-immunolabeled sections were analyzed on 40× images. Reporter-positive cells were identified using Smart Segmentation, and NeuN colocalization was assessed by overlap of reporter-positive somatic signal with NeuN immunoreactivity. NeuN colocalization was calculated as the proportion of reporter-positive cells exhibiting overlap with NeuN signal. This validation analysis was performed in a limited subset of AGS using a minimum of 2 NeuN-immunolabeled sections per reporter condition.

### Statistical analysis

Statistical analyses were performed using GraphPad Prism (version 10.6.1; GraphPad Software, San Diego, CA, USA). Data are presented as mean ± SEM unless otherwise noted. For quantitative analyses, values were averaged within each animal prior to statistical testing, and each animal was considered an independent biological replicate. Sample sizes (*n*) and statistical tests used for each comparison are reported in the corresponding figure legends. Groups with fewer than three animals were excluded from statistical analysis. A significance threshold of *p* < 0.05 was used for all analyses.

## Results

### AAV serotypes produce robust reporter expression with substantial neuronal enrichment in the AGS hypothalamus

Robust and anatomically localized reporter expression was observed for all tested AAV serotypes within the hypothalamus. Expression was confined to the injected hemisphere and, as intended, extended across the broader region surrounding the dorsomedial hypothalamus (DMH), spanning multiple coronal levels and adjacent hypothalamic areas (Fig. 1A–C). Higher-magnification images (Fig. 1D) show representative reporter-positive somata with overlapping NeuN immunoreactivity for both AAV1-EGFP and AAV-DJ-mCherry, supporting neuronal transduction in the AGS hypothalamus.

**Figure 1.**
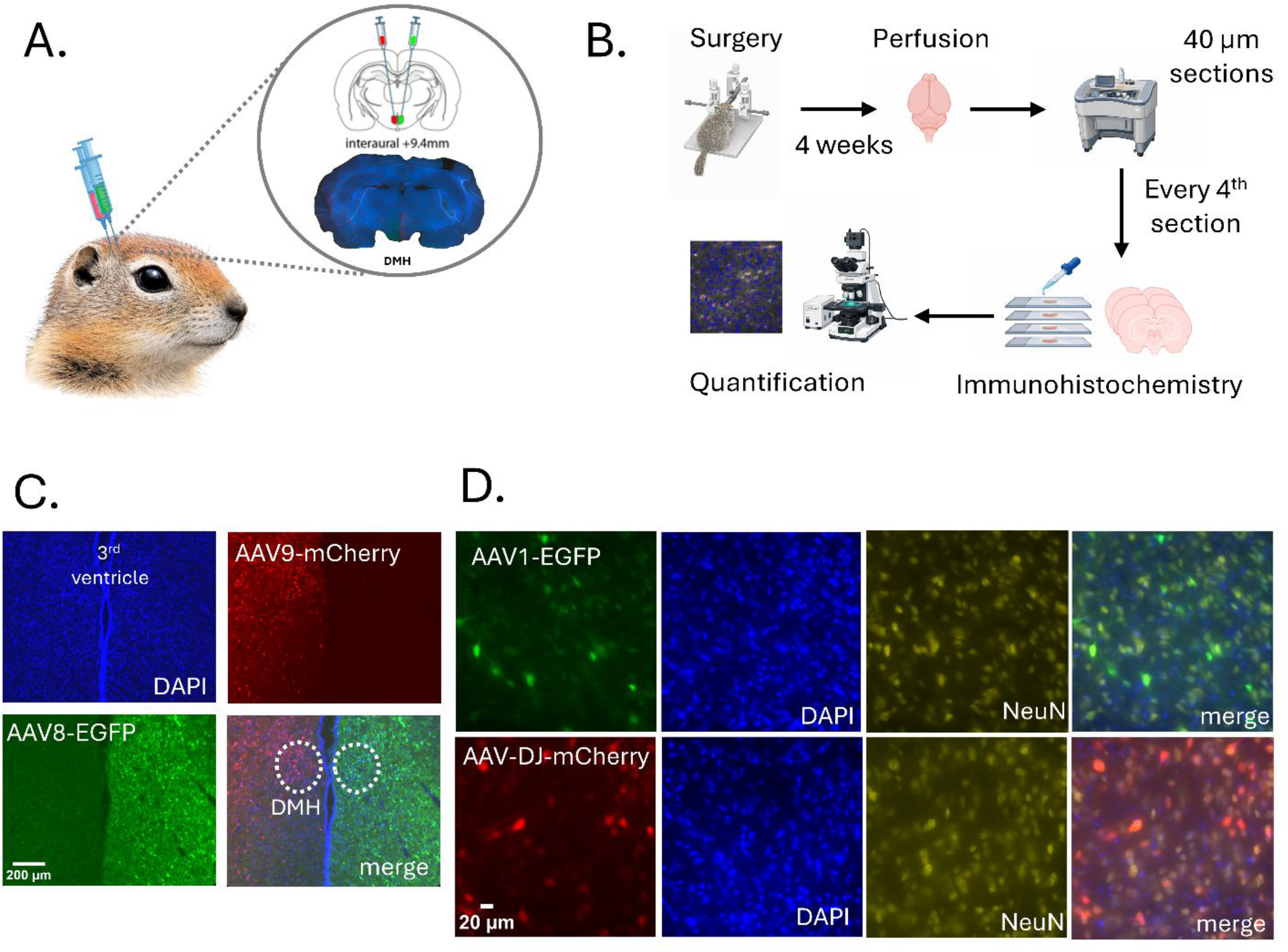
Experimental design, representative reporter expression, and neuronal colocalization in the AGS hypothalamus. (**A**)Schematic illustrating bilateral stereotaxic injection of adeno-associated viruses (AAVs) into the DMH of the AGS. Distinct fluorophore-expressing AAVs were delivered contralaterally. The injection site is shown relative to anatomical landmarks (interaural +9.4 mm). (**B**)Experimental workflow. Following stereotaxic surgery, animals recovered for 4 weeks before perfusion with 4% formalin. Brains were cryosectioned coronally at 40 µm thickness, and every fourth section was selected for immunohistochemical processing and quantification. (**C**)Representative low-magnification (4×) fluorescence images showing reporter expression within the hypothalamus. Individual channels for DAPI (blue), AAV9-mCherry (red), and AAV8-EGFP (green), as well as a merged image, are shown. Scale bar, 200 µm. (**D**)Representative high-magnification (40×) fluorescence images demonstrating reporter expression and colocalization with the neuronal marker NeuN within the DMH. Top row, AAV1-EGFP; bottom row, AAV-DJ-mCherry. Individual reporter, DAPI (blue), NeuN (yellow), and merged images are shown. Scale bar, 20 µm.

### AAV-DJ exhibits reduced cellular transduction rates relative to AAV1 in the AGS hypothalamus

To assess serotype-dependent differences in AAV performance in the AGS hypothalamus, we quantified transduction rates across multiple coronal sections and imaging fields using a standardized imaging and analysis pipeline. Cellular transduction rates were defined as the proportion of DAPI-positive nuclei exhibiting colocalization with reporter signal. Within-animal designs were used to enable direct serotype comparisons while controlling for inter-animal variability.

Comparison of AAV-DJ and AAV1 revealed a consistent serotype-dependent difference in transduction rates (Fig. 2A). Across animals and matched coronal sections, AAV-DJ exhibited lower transduction rates relative to AAV1. Mean cellular transduction rates were 17.2% ± 3.5% for AAV1 and 8.4% ± 2.9% for AAV-DJ. At the animal level, paired comparison of mean cellular transduction rates demonstrated a significant difference favoring AAV1 (paired t-test, *p* = 0.032; mean difference [AAV-DJ − AAV1] = −8.8%; 95% CI, −16.2% to −1.4%).

**Figure 2.**
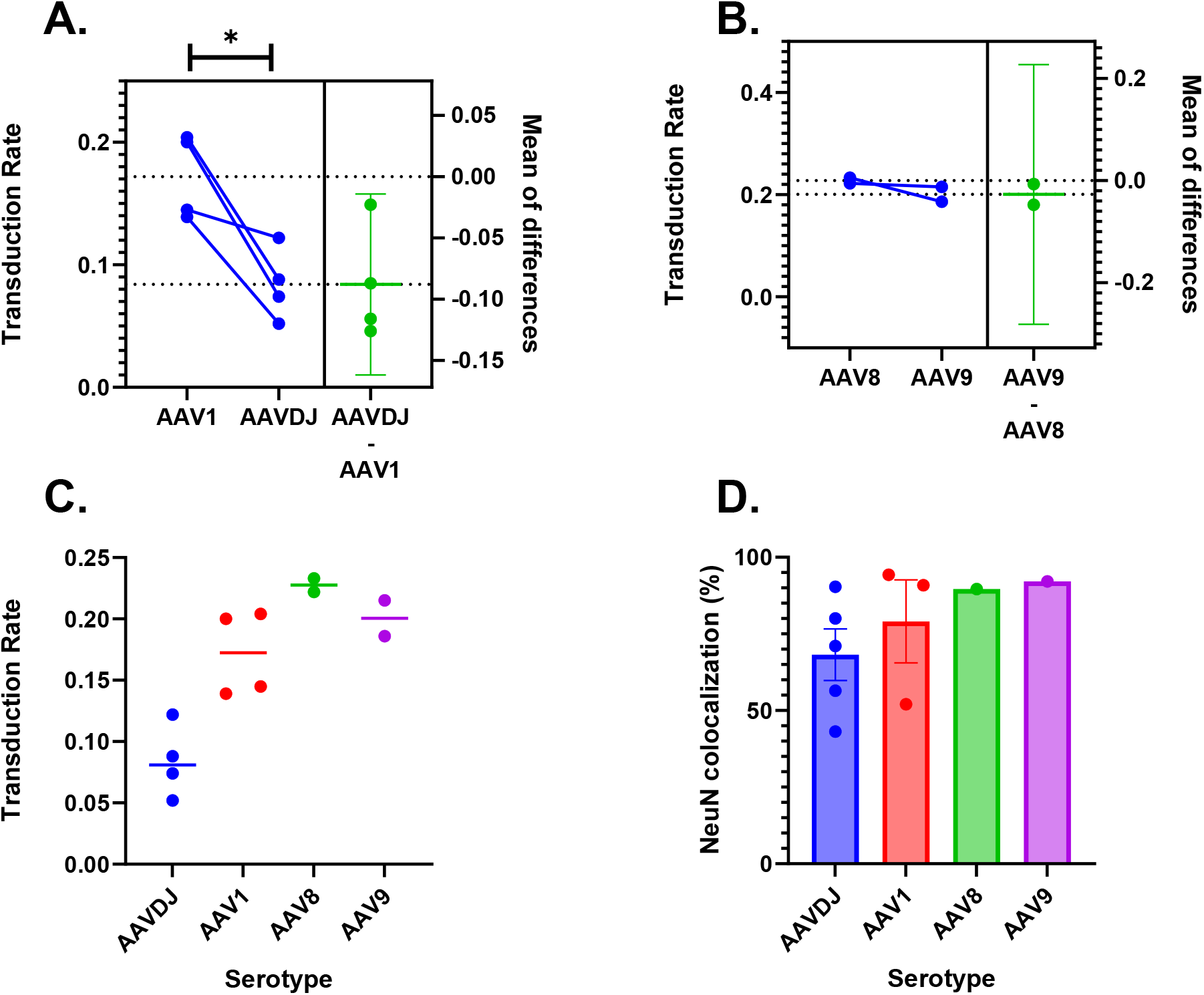
Serotype-dependent differences in transduction rates and NeuN colocalization in the AGS hypothalamus. **(A)** Within-animal comparison of AAV1 and AAV-DJ transduction rates across matched coronal sections of the hypothalamus. Left, paired animal-level mean transduction rates for AAV1 and AAV-DJ; lines connect values from the same animal (n = 4 animals). Right, estimation plot showing the mean paired difference in transduction rates (AAV-DJ − AAV1) with 95% confidence interval. Negative values indicate lower transduction rates for AAV-DJ relative to AAV1. A paired t-test identified a significant difference favoring AAV1 (*p = 0.032). **(B)** Within-animal comparison of AAV8 and AAV9 transduction rates across matched coronal sections of the hypothalamus. Left, paired animal-level mean transduction rates for AAV8 and AAV9; lines connect values from the same animal (n = 2 animals). Right, estimation plot showing the mean paired difference in transduction rates (AAV9 − AAV8). Transduction rates for AAV8 and AAV9 were similar, with the mean difference centered near zero. **(C)** Animal-level mean transduction rates across AAV serotypes. Each point represents one animal and horizontal lines indicate group means. AAV-DJ showed lower transduction rates than AAV1, whereas AAV8 and AAV9 overlapped within the limited animals analyzed. **(D)** NeuN colocalization of reporter-positive cells across AAV serotypes, assessed as the percentage of reporter-positive cells co-localized with NeuN. Each point represents the mean value from one animal, averaged across analyzed sections; bars indicate group means. NeuN colocalization was assessed in a limited subset of animals (AAV-DJ, n = 5; AAV1, n = 3; AAV8, n = 1; AAV9, n = 1), and these data are presented descriptively. AAV8 and AAV9 are shown descriptively due to the limited sample size.

Although confidence intervals remain relatively wide due to modest animal numbers, the directionality of the effect was consistent across animals, supporting a reproducible reduction in AAV-DJ transduction rates under these conditions.

### AAV8 and AAV9 show overlapping cellular transduction rates with no consistent serotype advantage

We next compared cellular transduction rates between AAV8 and AAV9 in the AGS hypothalamus (Fig. 2B). Mean cellular transduction rates were 22.8% ± 0.6% for AAV8 and 20.1% ± 1.5% for AAV9 (n = 2 animals). Because only two animals were available for this comparison, formal statistical testing was not performed. Under the experimental conditions tested, AAV8 and AAV9 exhibited overlapping cellular transduction rates, and no clear serotype advantage was observed.

### Cross-serotype summary highlights reduced cellular transduction rates of AAV-DJ

To place individual serotype comparisons in a broader context, we summarized animal-level mean transduction rates across all tested serotypes (AAV-DJ, AAV1, AAV8, and AAV9; Fig. 2C). Across experiments, AAV-DJ consistently occupied the lower range of observed transduction rates, whereas AAV1, AAV8, and AAV9 exhibited overlapping distributions.

Although direct statistical comparison across all four serotypes was limited by animal number and experimental pairing, the convergence of results across independent within-animal comparisons suggests that reduced transduction by AAV-DJ is a reproducible feature in this dataset. In contrast, differences among AAV1, AAV8, and AAV9 were comparatively small.

### Reporter expression shows substantial neuronal enrichment across serotypes

Given the use of the human synapsin (hSyn) promoter, which is expected to favor neuronal expression, we assessed neuronal enrichment by quantifying the proportion of reporter-positive cells that colocalized with the neuronal marker NeuN (reporter+NeuN+ / total reporter+ cells; Fig. 2D). Across serotypes, the majority of reporter-positive cells colocalized with NeuN, supporting substantial neuronal enrichment in the AGS hypothalamus. Mean NeuN colocalization was 76.8% ± 16.6% for AAV1 and 65.9% ± 21.2% for AAV-DJ. Descriptively, NeuN colocalization was 89.6% and 92.1% for AAV8 and AAV9, respectively. Although variability was greater for AAV1 and AAV-DJ, all serotypes demonstrated substantial neuronal enrichment, consistent with the expected bias of the hSyn promoter toward neuronal expression.

## Discussion

We compared the cellular transduction rates of AAV serotypes 1, 8, 9, and DJ in the hypothalamus of AGS and found that AAV1 produced significantly greater cellular transduction rates than AAV-DJ. In contrast, AAV8 and AAV9 exhibited cellular transduction rates comparable to AAV1. Cellular transduction rates were quantified by overlap of reporter signal (mCherry or EGFP) with DAPI-labeled nuclei. Across serotypes, reporter expression was also substantially enriched in NeuN-positive cells, supporting predominantly neuronal transduction under the hSyn promoter.

AAV1, AAV8, and AAV9 are naturally occurring serotypes, whereas AAV-DJ is an engineered hybrid capsid generated by DNA shuffling of multiple parental AAV serotypes to enhance transduction efficiency (Grimm et al. 2008). AAV-DJ was originally developed to improve upon limitations observed with earlier serotypes, including relatively inefficient transduction by AAV2 in certain tissues (Grimm et al. 2008). Within the CNS, AAV-DJ has also been reported to exhibit broader cellular tropism, including transduction of both neurons and astrocytes, rather than uniformly superior neuronal transduction. This context dependence may help explain its reduced cellular transduction rate in the AGS hypothalamus relative to AAV1. Direct comparisons between AAV-DJ and AAV1 in neuronal systems have revealed tissue-specific differences in transduction rates, including studies of spiral ganglion neurons in which AAV-DJ did not consistently outperform AAV1 (Lin, Nguyen, and Shibata 2025). Conversely, studies in mouse brain have reported enhanced neuronal transduction with AAV-DJ relative to AAV1; however, our findings in the AGS hypothalamus indicate that this relationship does not generalize across species or brain regions (Lunev et al. 2022).

Prior comparative studies in mouse brain have shown that AAV1 supports efficient neuronal transduction, with relative performance varying by brain region and experimental context (Aschauer et al. 2013). The relatively high cellular transduction rates observed for AAV1 in the AGS hypothalamus are therefore consistent with its established neuronal tropism. In contrast to engineered capsids such as AAV-DJ, whose performance may depend strongly on optimization within specific tissue contexts, AAV1’s naturally evolved capsid may confer advantages for neuronal entry or intracellular trafficking in the AGS hypothalamus, highlighting the importance of evaluating AAV tropism directly in non-model species.

The overlapping cellular transduction rates observed for AAV8 and AAV9 in the AGS hypothalamus may reflect shared neuronal tropism of these capsids, as similar performance of AAV8 and AAV9 has been reported in mammalian CNS tissue (Issa et al. 2023; Masamizu et al. 2011). Alternatively, high baseline cellular transduction rates within this region may limit the dynamic range available to detect capsid-dependent differences. Finally, the modest animal number reduced statistical power to resolve small effect sizes between these serotypes.

An important limitation of the present study is that contralateral injections preclude the determination of whether different serotypes transduce identical neuronal populations. It is possible that distinct serotypes target partially overlapping or non-overlapping subpopulations within the hypothalamus. Thus, comparable transduction rates across serotypes do not necessarily imply targeting of identical cellular ensembles. Another limitation is that warm ambient temperature may not fully deter expression of torpor. Short (∼24h) bouts with body temperatures reaching near ambient are observed at warm ambient temperature and were not monitored in this study (Olson et al. 2013).

Together, these findings provide an empirical framework for rational AAV serotype selection in the AGS, a non-model species with unique neurophysiological adaptations. Because capsid-dependent transduction performance can vary substantially across species and brain regions, direct validation of vector performance is essential prior to functional manipulation. The identification of serotypes with reliable hypothalamic transduction enables future studies targeting defined neuronal populations to investigate mechanisms underlying metabolic plasticity, metabolic suppression, neuronal plasticity, resistance to I/R injury and thermogenesis in this model. Reliable transduction of hypothalamic neurons enables future use of AAV-delivered sensors, chemogenetic, optogenetic, or gene-modulation approaches to interrogate neurochemical mechanisms and pathways regulating seasonal metabolic adaptations of the hibernating brain.

## Abbreviations

ABS: antibody dilution solution
AGS: Arctic ground squirrel
AAV: adeno-associated virus
DAPI: 4′,6-diamidino-2-phenylindole
DMH: dorsomedial hypothalamus
EGFP: enhanced green fluorescent protein
GC: genome copies
hSyn: human synapsin
I/R: ischemia/reperfusion
PBS: phosphate-buffered saline
SEM: standard error of the mean

## Acknowledgement

This work was supported by the National Institute of General Medical Sciences under award P20GM130443. L.E.F. is supported by the Clayton Foundation for Research, the W.M. Keck Foundation, the BD2 Foundation, the WoodNext Foundation, the National Science Foundation (award 24323797), and the Waggoner Center for Alcohol and Addiction Research. D.T. is supported by the Kuni Foundation Discovery Grants for Cancer Research. The authors thank Kelly Drew for helpful comments and feedback on the manuscript.

## Notes

### Competing Interest Statement

The authors have declared no competing interest.

